# Allele specific PCR for a major marker of levamisole resistance in *Haemonchus contortus*

**DOI:** 10.1101/2022.04.08.487639

**Authors:** Alistair Antonopoulos, Stephen R. Doyle, David J. Bartley, Alison A. Morrison, Ray Kaplan, Sue Howell, Cedric Neveu, Valentina Busin, Eileen Devaney, Roz Laing

## Abstract

*Haemonchus contortus* is a haematophagous parasitic nematode that infects small ruminants and causes significant animal health concerns and economic losses within the livestock industry on a global scale. Treatment primarily depends on broad-spectrum anthelmintics, however, resistance is established or rapidly emerging against all major drug classes. Levamisole (LEV) remains an important treatment option for parasite control, as resistance to LEV is less prevalent than to members of other major classes of anthelmintics. LEV is an acetylcholine receptor (AChR) agonist that, when bound, results in paralysis of the worm. Numerous studies implicated the AChR sub-unit, ACR-8, in LEV sensitivity and in particular, the presence of a truncated *acr-8* transcript or a deletion in the *acr-8* locus in some resistant isolates. Recently, a single non-synonymous SNP in *acr-8* conferring a serine-to-threonine substitution (S168T) was identified that was strongly associated with LEV resistance. Here, we investigate the role of genetic variation at the *acr-8* locus in a controlled genetic cross between the LEV susceptible MHco3(ISE) and LEV resistant MHco18(UGA 2004) isolates of *H. contortus*. Using single worm PCR assays, we found that the presence of S168T was strongly associated with LEV resistance in the parental isolates and F3 progeny of the genetic cross surviving LEV treatment. We developed and optimised an allele-specific PCR assay for the detection of S168T and validated the assay using laboratory isolates and field samples that were phenotyped for LEV resistance. In the LEV-resistant field population, a high proportion (>75%) of L_3_ encoded the S168T variant, whereas the variant was absent in the susceptible isolates studied. These data further support the potential role of *acr-8* S168T in LEV resistance, with the allele-specific PCR providing an important step towards establishing a sensitive molecular diagnostic test for LEV resistance.

## 1. Introduction

*Haemonchus contortus* is one of the most pathogenic and economically important gastrointestinal nematodes (GIN) of small ruminants, and is responsible for significant welfare issues and production losses for the sheep sector worldwide (Miller et al., 2012; Emery et al., 2016; Kotze et al., 2016; Besier et al., 2016; Sallé et al., 2019). Control of haemonchosis relies heavily on prophylactic and therapeutic use of broad-spectrum anthelmintics (Kotze et al., 2016), however, *H. contortus* shows a remarkable capacity to quickly develop resistance under drug pressure (Kaplan, 2020). Resistance to four of the major classes of anthelmintic (benzimidazoles, imidazothiazoles such as levamisole (LEV), macrocyclic lactones and amino-acetonitrile derivatives) has been reported (Sangster and Gill, 1999; Wolstenholme et al., 2004; Gilleard, 2013; Van den Brom et al., 2015) and the prevalence of multidrug resistance is increasing in many areas (Kaplan, 2004; Geurden et al., 2014; Kotze et al., 2016). Resistance to LEV is less common than to other broad-spectrum anthelmintics (Cernanská et al., 2006; Van den Brom et al., 2013; Crook et al., 2016; Rose-Vineer et al., 2020) and, as such, it remains an important control option for the livestock industry.

LEV is a selective agonist of nematode acetylcholine receptors (AChRs) (Aceves et al., 1970; Martin, 1997; Kopp et al., 2009; Martin et al., 2012). Nematode AChRs are pentameric ligand-gated ion channels composed of five identical (homomeric) or related (heteromeric) subunits (Duguet et al., 2016). The LEV sensitive AChR in *H. contortus* is a heteropentameric complex formed from subunits UNC-38, UNC-29, UNC-63, and ACR-8 with one subunit present twice (Boulin et al., 2011; Blanchard et al., 2018). The canonical binding site for acetylcholine and the allosteric binding site for LEV are thought to be distinct but located at the interface between two receptor subunits, with ACR-8 and UNC-63 likely candidates among others (Martin et al., 2012; Duguet et al., 2016). Multiple lines of evidence, including functional reconstitution and RNAi experiments, have shown that ACR-8 plays a key role in conferring LEV sensitivity to the *H. contortus* AChR (Boulin et al., 2011; Blanchard et al., 2018).

Current understanding of LEV resistance in *H. contortus* has largely focused on three putative mechanisms: (i) the expression of truncated transcripts of *acr-8* and *unc-63* (termed *acr-8b* and *unc-63b* respectively) (Neveu et al., 2010; Fauvin et al., 2010; Boulin et al., 2011; Barrère et al., 2014), (ii) altered expression patterns of AChR subunits (including *acr-8* and *unc-63*) (Sarai et al., 2013; Sarai et al., 2014; Raza et al., 2016), and (iii) a variable length deletion present in the second intron of the *acr-8* gene (Williamson et al., 2011; Barrère et al., 2014; dos Santos et al., 2019). The deletion in intron 2 has also been correlated with expression of *acr-8b* (Barrère et al., 2014), and it has been proposed that the two could be involved in LEV resistance (Barrère et al., 2014; dos Santos et al., 2019).

Diagnosis of anthelmintic resistance, including LEV resistance, typically relies on the faecal egg count reduction test (FECRT), which compares egg counts between pre- and post-drug anthelmintic administration (Coles et al., 1992). While faecal egg counts are relatively simple to perform and are useful for general flock management, the FECRT has significant drawbacks including poor sensitivity for detection of resistance; for the BZs, at least 25% of the population had to be resistant for detection (Martin et al., 1989). This presents serious concerns for managing the emergence of resistance, particularly for drugs where clinical efficacy remains relatively high such as LEV (Rose-Vineer et al., 2021).

Molecular diagnostic tests have been demonstrated for multiple benzimidazole (BZ) resistance SNPs in parasitic nematodes using various methodologies, including allele specific (AS)-PCR, restriction fragment length polymorphism (RFLP), and pyrosequencing (Winterrowd et al., 2003; Tiwari et al., 2006; von Samson-Himmelstjerna et al., 2009; Mohanraj et al., 2017). A recent development in large scale surveillance is the ‘Nemabiome’ approach, which makes use of deep amplicon sequencing of barcoded PCR products (Avramenko et al., 2015). Although originally developed for species identification and quantification, it was recently adapted to assess the presence of BZ resistance SNPs by deep sequencing of β–tubulin amplicons (Avramenko et al., 2019; Melville et al., 2020). However, to date, no molecular diagnostic assays are described for LEV resistance surveillance.

We recently used bulk segregant analysis of a genetic cross between LEV resistant and LEV susceptible isolates of *H. contortus* together with whole-genome sequencing to identify a discrete region on Chromosome V under LEV selection (Doyle et al., 2022). This region contained gene HCON_00151270, the *H. contortus* orthologue of *C. elegans acr-8*. We identified a single non-synonymous SNP in *H. contortus acr-8* exon 4 encoding a serine-to-threonine substitution (PRJEB506: 31,521,884 bp; GCT-GGT) and refer to this variant as S168T. This variant was highly differentiated between the susceptible and resistant parental isolates, and its frequency increased significantly in the progeny of the genetic cross after LEV selection. Notably, the serine residue at this position was highly conserved amongst parasitic Clade V nematodes, yet a serine-to-threonine substitution was also found at the corresponding position of the *acr-8* orthologue in LEV resistant *Teladorsagia circumcincta* (Choi et al., 2017; Doyle et al., 2022). Here we build upon these population-level findings to validate the S168T variant using single worm genotyping of *H. contortus* L_3_ from both parental isolates and the genetic cross pre- and post-LEV treatment, and describe the development of an optimised allele-specific PCR (AS-PCR) to detect LEV resistant and susceptible *acr-8* alleles in laboratory and field populations.

## 2. Materials and Methods

### 2.1 Animal handling and ethics statement

All experimental procedures were examined and approved by the Animal Welfare Ethical Review Board of the Moredun Research Institute, Penicuik, Scotland and were conducted under approved UK Home Office licenses following the Animals (Scientific Procedures) Act of 1986. The Home Office licence number is PPL 60/03899.

### 2.2 *H. contortus* isolates

The *H. contortus* samples used in this study were from the LEV susceptible MHco3(ISE) (Otsen et al., 2001; Roos et al., 2004) isolate, the LEV resistant MHco18(UGA2004) (Williamson et al., 2011) isolate, and the progeny of genetic cross between these isolates termed MHco3/18, originally described in Doyle et al. (2018). In addition, two LEV susceptible geographically divergent isolates: MHco4(WRS) (Van Wyk et al., 1989) and MHco10(CAVR) (Le Jambre, 1993) were also used. All isolates were maintained and passaged through parasite-naive sheep at the Moredun Research Institute. In brief, the MHco3/18 line was established by crossing immature MHco3(ISE) females with immature MHco18(UGA2004) males at 14 days post infection. Infective larvae from the MHco3/18 cross were used to infect parasite naïve lambs to generate two further filial generations, followed by *in vivo* selection of the F2 generation by treatment of three infected animals with 7.5 mg/kg body weight levamisole hydrochloride (Levacide, Low Volume, Norbrook Laboratories Ltd) (Doyle et al., 2022). Eggs from the F3 generation were collected from donor sheep pre-treatment and again at 21 days post-treatment, after which they were cultured to L_3_ (Coop et al., 1982). All populations of L_3_ were either maintained at 8°C in tap water or cryopreserved in liquid nitrogen. Adult worms collected from sheep at post-mortem were sexed then cryopreserved in liquid nitrogen.

Field samples were collected in the Southern USA and phenotyped using a commercial larval development assay (DrenchRite®). For LEV, resistant populations have an EC50 >= 1.56 μM and susceptible populations have an EC_50_ <=0.78 μM. Farm 001 was highly LEV resistant (EC_50_: 9.36 μM) and Farm 002 was LEV susceptible (EC_50_: 0.57 μM). Farm 001 corresponds to Farm 7 in Doyle et al. (2022), Farm 002 corresponds to Farm 4 in Doyle et al. (2022).

### 2.3 DNA extraction

Pooled gDNA was extracted from frozen aliquots of ~10,000 L_3_ by a modified phenol chloroform extraction. Briefly, 20 μl (20 mg/μl) Proteinase K (Invitrogen, 25530015) and 300 μl lysis buffer (200 mM NaCl, 100 mM Tris-HCl, 30 mM EDTA, 0.5% SDS) was added to a pellet of worms, incubated at 55°C for 2 hr, after which 10 μl (10 mg/ml) RNase A was added and incubated at 37°C for 10 min. Phenol/chloroform/isoamyl alcohol (25:24:1; 550 μl) was then added, shaken, and incubated at R/T for 5 min, then centrifuged at 14000 × g for 15 min at R/T. The top layer was removed to a fresh tube and 200 μl chloroform was added, and the resultant mixture centrifuged at 14000 × g for 15 min at R/T. The top layer was again removed to a fresh tube and combined with 0.1 volume of 3 M sodium acetate, three volumes of 100% ethanol, and 2 μl glycogen, followed by overnight incubation at −80°C. The following day, the mixture was centrifuged at 14000 × g at 4°C for 45 min, the supernatant aspirated and 500 μl 70% ethanol added before centrifuging at 14000 × g at 4°C for 5 min. Finally, the pellet was air dried, and resuspended in 10 μl elution buffer (QIAgen, 19086). The DNA concentration was calculated using a Qubit Spectrophotometer (Qubit dsDNA HS kit Invitrogen, Q33231) and diluted to 100 ng/μl.

Crude lysates of single worms were produced as follows: 10 μl lysis buffer (100X stock solution: 1000 μl Direct PCR Lysis Reagent (Cell) (Viagen, 301-C), 50 μl 1 M DTT (Invitrogen, P2325), and 10 μl Proteinase K (100 mg/ml) (Invitrogen, 25530049) was aliquoted per well of a 96 well PCR plate. One L_3_ was picked into each well in a volume of <1 μl of water. Lysates were incubated at 60°C for 2 hr, followed by 85°C for 45 min to denature the Proteinase K.

### 2.4 Primer design and amplicon sequencing

Illumina reads from whole-genome sequencing of pools of 200 MHco3(ISE), MHco18(UGA2004), and MHco3/18 L_3_ (Doyle et al., 2022) were mapped to the PRJEB506 MHco3(ISE)_4.0 assembly (Doyle et al., 2020). Reads aligning to the *acr-8* genomic locus (HCON_00151270, 2020-04-PREB506.WBPS13 Wormbase annotation) were visualised in Geneious (Biomatters Ltd: 11.1.5) to identify between-isolate polymorphisms. Consensus sequences were generated for each isolate to simplify primer design. However, population-level data was also examined to ensure that there were no moderate frequency mutations that could disrupt primer binding sites. Generic and AS-PCR primers (Table 1) were manually designed in Geneious and supplied by Eurofins Genomics, Ebersberg, Germany. Deletion spanning primers (Hco-indel-F and Hco-indel-R) were designed ~50 bp up and downstream of the deletion locus, to allow sufficient size difference between amplicons for discrimination of PCR products via gel electrophoresis (Figure 1A). Indel specific primer Hco-Indel-Ins-R was designed within the region of the deletion locus that was conserved in the 97 bp deletion described by Doyle et al. (2022) and the 63 bp deletion described by Barrere et al. (2014) (Figure 1B). This was done to account for potential variation between isolates in the size and locus of the deletion in individual L_3_ based on previous reports (Barrère et al., 2014).

**Table 1:**
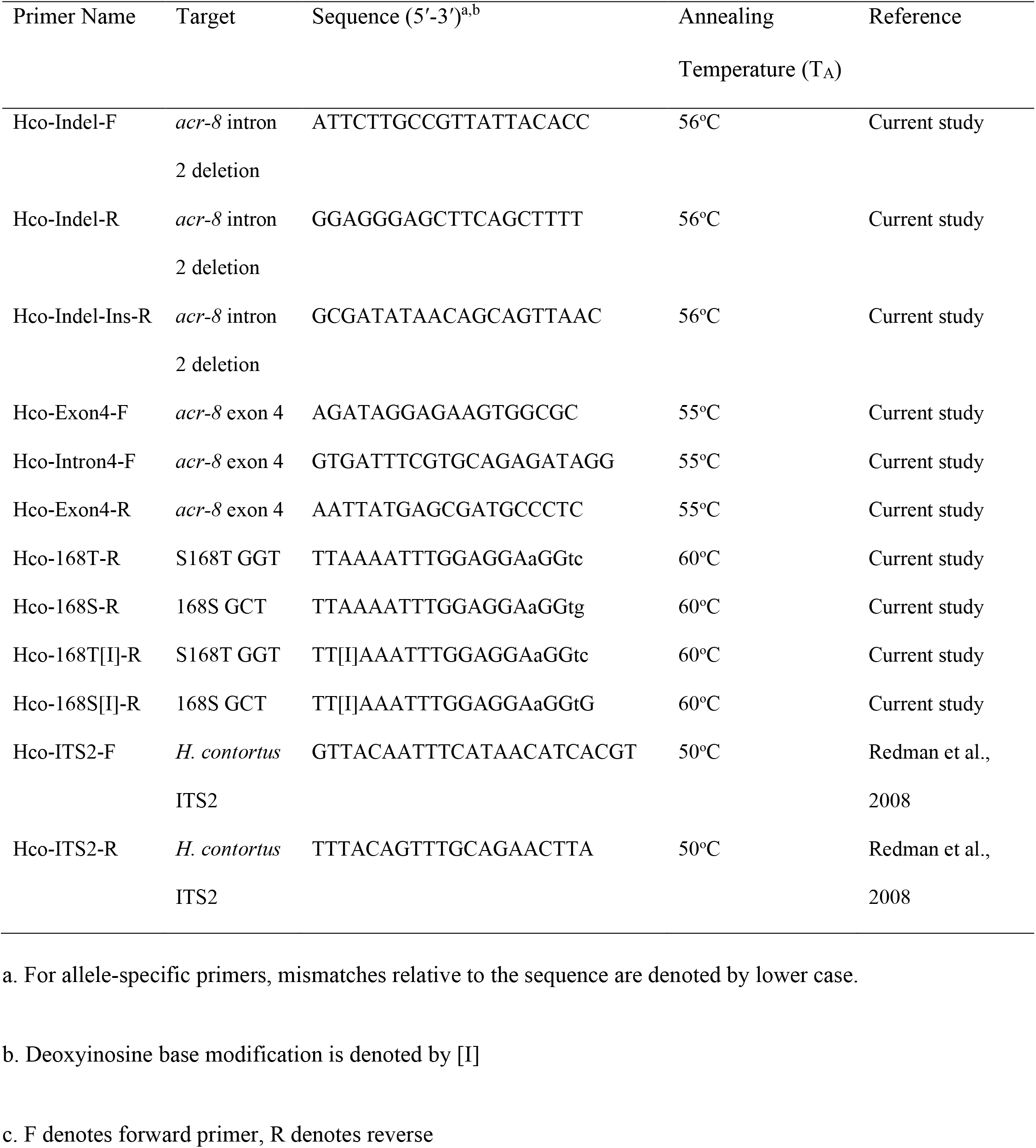
Table of primer sequences used in this study.

**Figure 1.**
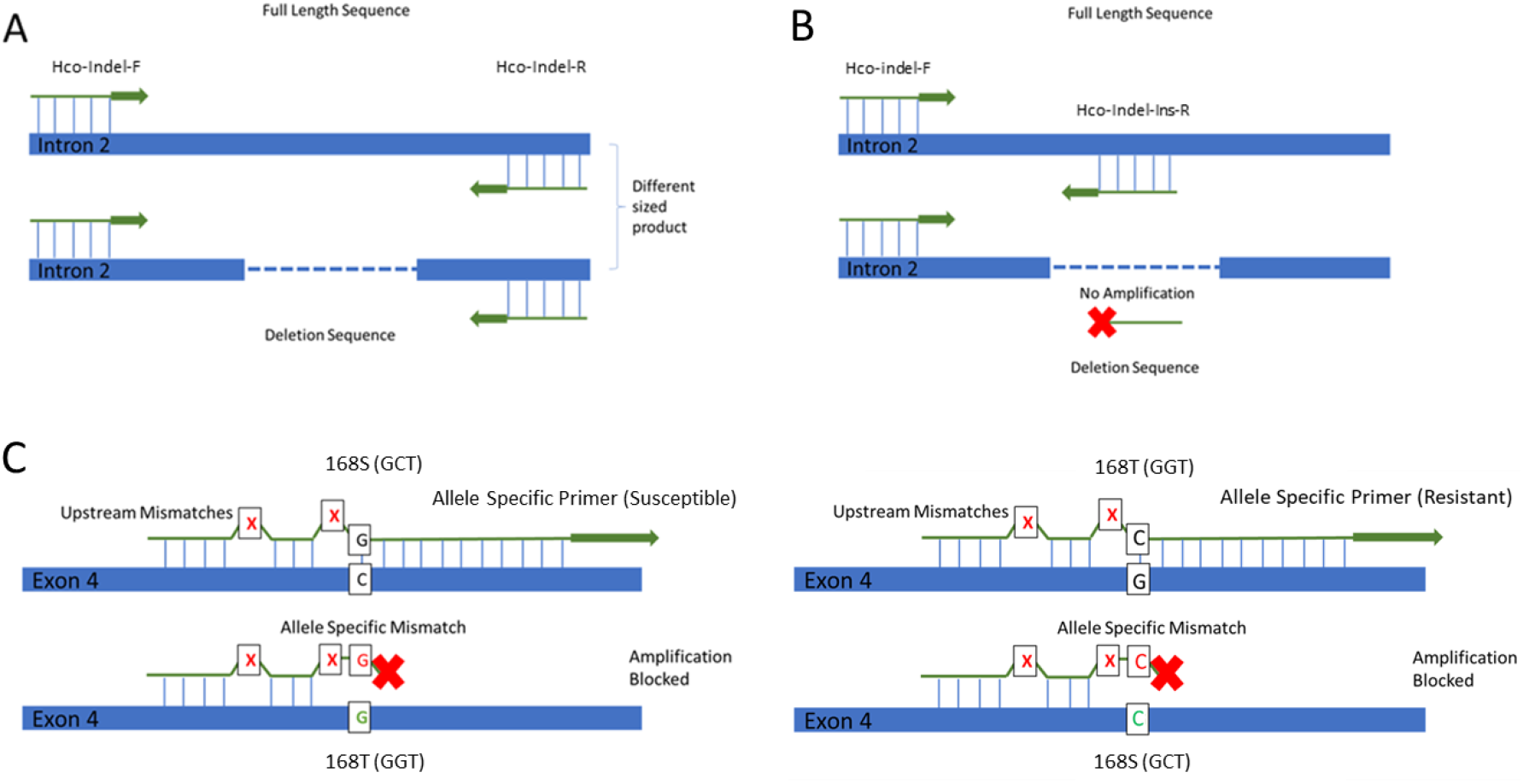
Schematic representation of primer design. A: Indel spanning primers (Table 1: Hco-Indel-R and Hco-Indel-F). If the deletion is present, then the product size will differ from the full-length allele, allowing for a size discrimination PCR to determine the genotype of the individual. B: Deletion specific primer (Table 1: Hco-Indel-Ins-R). The deletion specific primer lies wholly within the deleted sequence, thus blocking amplification if the deletion is present. C: Schematic representation of allele-specific primer function. The allele-specific primer binds upstream of the discriminating allele in both sensitive and resistant DNA, however, the ability of the primer to facilitate DNA extension during PCR depends specifically on the nucleotide at the 3’ end of the primer. Additional nucleotide mismatches at the 2^nd^ and 5^th^ position from the 3’ end of the primer (indicated by red X) are identical in each set and enhance the destabilisation of the polymerisation complex in the case of an allele specific mismatch at the 3’ position to discriminate between the two alleles GGT (168T/resistant) and GCT (168S/susceptible). The 3’ mismatch is allele specific, and as shown by the example above will prevent polymerisation and amplification if the S168T variant is present. In the resistant primer assay, the opposite occurs.

AS-PCR primers were designed with mismatches introduced at the 2^nd^ and 5^th^ bases from the 3′ end, for both the susceptible and resistant primers, in addition to a SNP-specific mismatch (PRJEB506 position: 31,521,884 bp) at the most 3′ base for the resistant primers only (Figure 1C). For two genetically divergent laboratory isolates (MHco4(WRS) and MHco10(CAVR)), published whole genome sequencing reads (Redman et al., 2012; Gilleard, 2013) were mapped to the genome and the *acr-8* genomic locus was visualised as described above. Two synonymous SNPs, C-T (PRJEB506 position: 31,521,901) and A-G (PRJEB506 position: 31,521,895), were identified in a high proportion of reads from both MHco4(WRS) and MHco10(CAVR). As both SNPs lie within the primer binding site for the allele-specific primers, deoxyinosine bases were substituted at positions 17, and 17 and 11 from the 5′ end of the Hco-168T-R and Hco-168S-R primers, respectively. These primers are termed Hco-168T[I]-R / Hco-168S[I]-R and Hco-168T[II]-R / Hco-168S[II]-R, where I denotes the addition of a deoxyinosine base.

### 2.5 Reaction-specific conditions for PCRs and RFLP

#### 2.5.1 *acr-8* intron 2 deletion PCR

The intron 2 region was amplified from genomic DNA with GoTaq G2 Flexi DNA polymerase (Promega, M7801) in a final volume of 20 μl as follows: 1 μl of crude lysate was added to 5 μl 5X Green GoTaq Flexi Buffer, 8 μl 25mM MgCl_2_, 1 μl 10 mM dNTP mix (Promega, U1511), 1 μl F and R primer (100 pmol), 0.25 μl GoTaq G2 Flexi DNA polymerase, 3.75 μl DNAse/RNAse-free water and amplified as follows: initial denaturation 95°C for 2 min, followed by 35 cycles (denaturation 95°C for 30 s, primer annealing temperature (T_A_) °C for 30 s, and extension 72°C for 15 s with a final extension at 72°C for 5 min) using primers Hco-Indel-F and R, or Hco-Indel-F and Hco-Indel-Ins-R. All PCR products were visualised on 2% agarose gel with SafeView Nucleic Acid Stain. Identity of PCR products was confirmed by capillary sequencing (Eurofins, Ebersberg, Germany).

#### 2.5.2 Two-step nested PCR and RFLP for S168T

The exon 4 region (PRJEB506 position: 31,521,758 – 31,522,025 bp) was amplified from single worm lysates as described in 2.5.1 except using primers Hco-Intron4-F and Hco-Exon4-R. 1 μl of the PCR product from round 1 was diluted 1:20 and added to 19 μl GoTaq G2 Flexi PCR reaction mix and amplified as follows: initial denaturation 95°C for 2 min, followed by 40 cycles (denaturation 95°C for 30 s, annealing T_A_ °C for 30 s, and extension 72°C for 15 s) with the final extension at 72°C for 5 min. 20 μl of nested PCR product was digested using *Ava*II (NEB, R0153S) for 7-12 h at 37°C in CutSmart buffer (NEB, B7204S). Digested products were then visualised on a 4% NuSieve (Lonza, 50090) agarose gel with SafeView Nucleic Acid Stain.

#### 2.5.3 Optimisation of AS-PCR for S168T with cloned fragments

Cloned *acr-8* exon 4 fragments for both S168T variants (GCT from MHco3(ISE), GGT from MHco18(UGA2004)) were used to optimise the AS-PCR; these provided abundant template and removed the complexity of additional polymorphisms at the primer binding sites in individual worms during early optimisation. Target sequences for *acr-8* exon 4 were amplified from MHco3(ISE) and MHco18(UGA2004) single worm lysates (2.3; 2.5.4), using Phusion HiFi PCR (Hco-Intron4-F/Hco-Exon4-R) and cloned using TOPO™ TA Cloning™ with One Shot™ TOP10 chemically competent *E. coli* cells (Invitrogen, K450001) or XL10-Gold ultracompetent *E. coli* cells (Agilent, 200315) according to the manufacturer’s instructions. Colonies were cultured on LB-Agar with 100 μg/ml ampicillin (Sigma Aldrich, A9518) and screened using X-gal (Promega, V3941) blue/white colony screening. White colonies were picked and further screened with RFLP (section 2.5.2) to identify the presence of GGT or GCT exon 4 fragments. Colonies containing the required insert/allele were sub-cultured on LB-Agar with 100 μg/ml ampicillin. Plasmid isolation was carried out using QIAprep Spin mini-prep kit (QIAgen, 27104) according to manufacturer instructions. Liquid cultures were grown up overnight in 5 ml of LB broth with 100 μg/ml ampicillin. Purified plasmid was capillary sequenced by Eurofins Genomics, Ebersberg, Germany according to company instructions.

Primer candidates Hco-168T-R and Hco-168S-R were predicted to discriminate between GCT and GGT *in silico* based on WGS for both isolates and sequencing of the clones. Gradient PCR using plasmid as template was used initially to establish the optimal T_A_ of each primer candidate and the optimal forward primer pairing. Best results were obtained by pairing Hco-Exon4-F/Hco-168T-R (132 bp amplicon) or Hco-Intron4-F/Hco-168S-R (148 bp amplicon). Touchdown-PCR cycles were then added to improve specificity and eliminate non-specific amplification. Extensive experimentation was carried out to optimise PCR conditions including polymerase, temperature, touchdown cycle number and duration, optimal dilution of first round PCR product, and dNTP and primer concentration.

#### 2.5.4 AS-PCR for S168T with single worm lysates

First-round PCR amplification of exon 4 was carried out on 1 μl of single worm lysate using Phusion Green High-Fidelity DNA Polymerase (Thermo Scientific, F534S) in a final volume of 20 μl as follows: 4 μl 5X HF reaction buffer, 0.2 μl 10 mM dNTP mix, 0.1 μl Hco-Intron4-F/Hco-Exon4-R primers (100 pmol), 0.2 μl Phusion DNA Polymerase, 14.5 μl DNAse/RNAse-free water. The PCR programme was as follows: initial denaturation for 30 s at 98°C, followed by 35 cycles (denaturation 98°C for 10 s, annealing T_A_ °C for 10 s, and extension 72°C for 10 s) with the final extension 72°C for 5 min.

A second-round PCR amplification with allele-specific primers was carried out using DreamTaq DNA Polymerase (Thermo Scientific, EP0702) PCR in a final volume of 20 μl as follows: 2 μl 10X reaction buffer, 0.05 μl 10 mM dNTP mix, 0.025 μl Hco-Exon4-F/Hco-168T-R or Hco-Intron4-F/Hco-168S-R (100 pmol), 0.2 μl DreamTaq polymerase, 16.9 μl DNAse/RNAse-free water, and 1 μl of the first-round PCR product diluted 1:10-1:20. The PCR programme using a touchdown PCR was as follows: initial denaturation for 2 min at 95°C, followed by 12 touchdown PCR cycles (denaturation 95°C for 30 s, annealing first temperature 68°C for 30 s, annealing last temperature 60°C (decreasing in equal increments per cycle) for 30 s, and extension 72°C for 15 s), followed by 10 PCR cycles (denaturation 95°C for 30 s, annealing temperature 60°C for 30 s, and extension 72°C for 15 s) with the final extension of 72°C for 5 min. PCR products were visualised on a 2% agarose gel stained with SYBR Safe DNA Gel Stain (Thermo Fisher, S33102).

#### 2.5.5 ITS2 *H. contortus* species identification PCR

ITS2 species identification PCR was carried out on 1 μl of single worm lysate using Phusion Green High-Fidelity DNA Polymerase (Thermo Scientific, F534S) in a final volume of 20 μl as described in 2.5.4 except using Hco-ITS2-F/Hco-ITS2-R (Redman et al., 2008). The PCR programme was as follows: initial denaturation for 30 s at 98°C, followed by 35 cycles (denaturation 98°C for 10 s, annealing T_A_ °C for 10 s, and extension 72°C for 10 s) with the final extension 72°C for 5 min.

## 3. Results

### 3.1 Detection of the *acr-8* intron 2 indel in LEV resistant and susceptible *H. contortus* L_3_

A previous study suggested an association between an indel in intron 2 of *acr-8* and LEV resistance in *H. contortus* (Barrère et al., 2014). Alignment of WGS to the reference genome showed a 97 bp deletion in MHco18(UGA2004) and MHco3/18 that increased in frequency from 73.47% to 86.58% in MHco3/18 F3 progeny following LEV selection of the F2 parental generation (Doyle et al., 2022). Primers were designed to bind ~50 bp upstream and downstream of this region and used to investigate its utility as a genetic marker of LEV resistance by size discrimination PCR. Genotyping of individual L_3_ from MHco18(UGA2004) and MHco3(ISE) demonstrated that the deletion was present in a high proportion of both populations (Figure 2A and B; Supplementary Table 1). Of 135 MHco3(ISE) worms genotyped, 64% (n=87) were found to be homozygous for the deletion compared with 92% (n=44) of the 48 MHco18(UGA2004). A confirmatory PCR with a primer designed to span the deletion locus (i.e., can only anneal if the deletion is absent) (Figure 2C) was performed. 100% concordance was found between the indel and confirmatory PCR (Figure 2C and D). Overall, these data show that the intron 2 deletion is present at a relatively high frequency in the MHco3(ISE) isolate of *H. contortus* (Figure 2B; Supplementary Figure 1).

**Figure 2:**
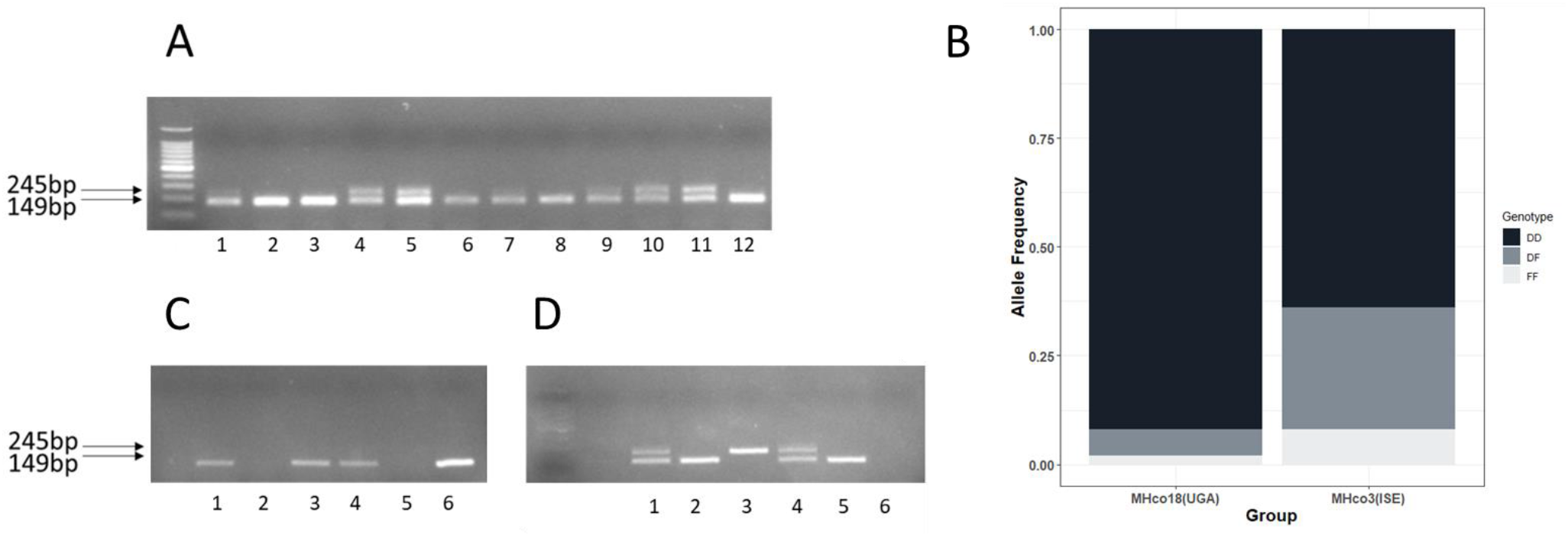
Summary of *acr-8* intron 2 deletion in MHco18(UGA2004) and MHco3(ISE) L_3_. Size discrimination PCR (Figure 1A) was developed to genotype individual L_3_ based on the presence or absence of the deletion allele. A deletion specific confirmation PCR (Figure 1B) was then developed to validate the results of the size discrimination PCR. A: Example gel of single worm PCR of MHco18(UGA2004) and MHco3(ISE) with indel spanning (~50 bp up and downstream of deletion locus) primer set (Hco-Indel-F/Hco-Indel-R). Odd numbers: MHco18(UGA2004). Even numbers: MHco3(ISE). 245 bp band corresponds to full length allele, 149 bp band corresponds to deletion allele. B: Bar chart showing proportion of deletion (D) allele detected by RFLP in LEV resistant MHco18(UGA2004), LEV susceptible MHco3(ISE), and MHco3/18 genetic cross. Y axis: genotype frequency. Genotypes DD: homozygous deletion; DF: heterozygous; FF: homozygous full length. C: Single worm PCR with indel internal (will only anneal when deleted sequence is present) confirmation (Hco-Indel-F/Hco-Indel-Ins-R) primer set: 1,3,5 MHco18(UGA2004); 2,4 MHco3(ISE) single L_3_ worms; 6 Positive Control (100 ng pooled L_3_ MHco3(ISE) gDNA. D: Single worm PCR with indel spanning (Hco-Indel-F/Hco-Indel-R) primer set: 1,3,5 MHco18(UGA2004), 2,4; MHco3(ISE) single L_3_, 6 NTC. The same individual L_3_ lysate was used for PCR shown in C and D.

### 3.2 Detection of S168T (GCT/GGT) in LEV resistant *H. contortus* L_3_

Having identified a high frequency of individuals with the *acr-8* indel in a fully LEV susceptible population, we focused our efforts on validating variant S168T in individual worms. Initially, we developed an RFLP assay, which could discriminate between the two alleles based on the introduction of an *Ava*II restriction site by the variant. This initial assay was useful for identifying GCT (encoding serine = susceptible/S) homozygotes and individuals with at least one GGT (encoding threonine = resistant/R) allele, and we were able to show that the S168T allele was present in a high proportion of MHco18(UGA2004) and MHco3/18, and absent in MHco3(ISE) (Figure 3A). However, it was not always possible to differentiate GGT homozygotes and heterozygotes, due to incomplete digestion (Figure 3B). For this reason, we focused our efforts on the development of an AS-PCR assay to discriminate between the two alleles (Figure 4).

**Figure 3:**
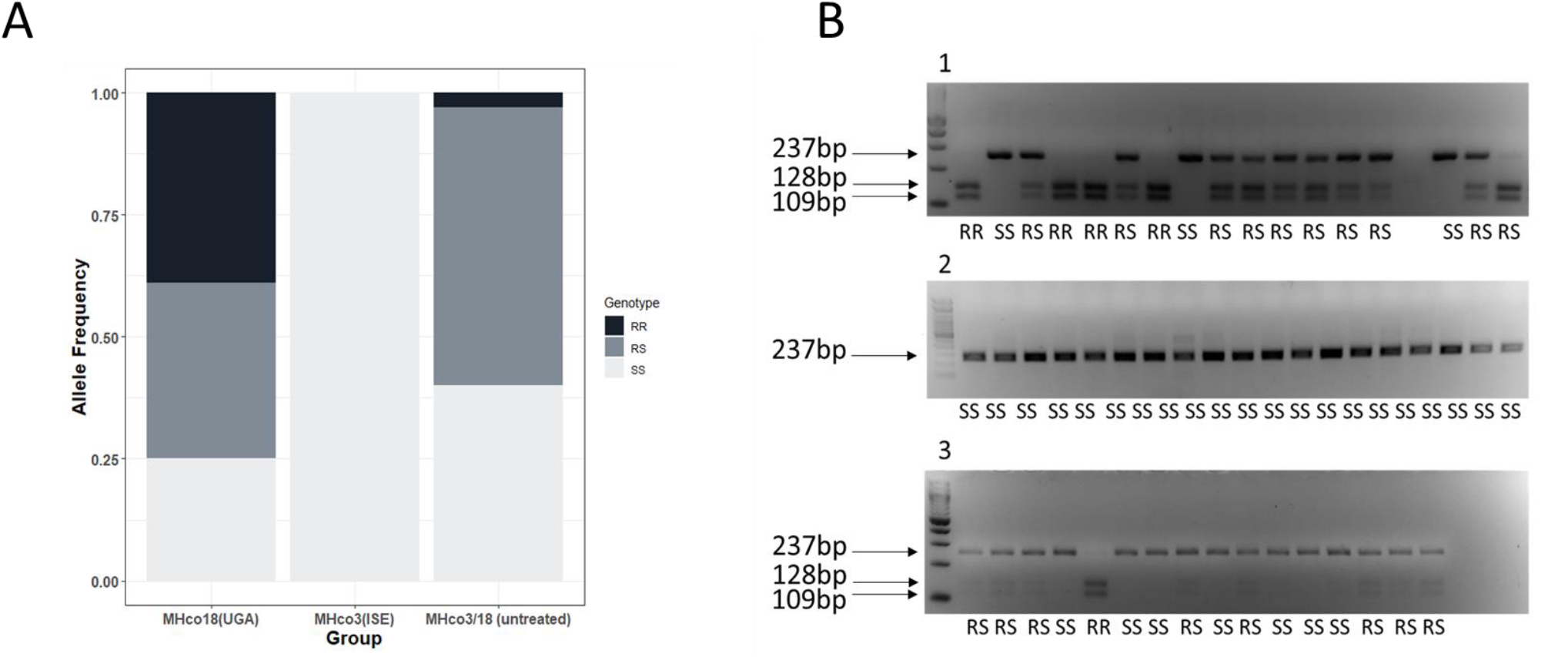
Summary of RFLP analysis of MHco18(UGA2004), MHco3(ISE) and MHco3/18 L_3_. S168T (GCT-GGT) mutation at codon 168 introduces an *Ava*II restriction site. Restriction Fragment Length Polymorphism (RFLP) was developed to genotype individual MHco3(ISE), MHco18(UGA2004), and MHco3/18 L_3_ for the presence of either the GCT (susceptible - S) or the GGT (resistant - R) allele. A: Bar chart showing proportion of GGT allele detected by RFLP in LEV resistant MHco18(UGA2004), LEV susceptible MHco3(ISE), and MHco3/18. Y axis: genotype frequency. B: Detection of GGT by single L_3_ *Ava*II RFLP in LEV resistant MHco18(UGA2004), LEV susceptible MHco3(ISE), and MHco3/18 genetic cross. 1: MHco18(UGA2004); 2: MHco3(ISE); 3: MHco3/18. 237 bp bands represents 168S GCT sequence. 128 bp and 109 bp bands represent GGT sequence. Predicted genotypes RR: homozygous GGT; RS: heterozygous; SS: homozygous GCT.

**Figure 4:**
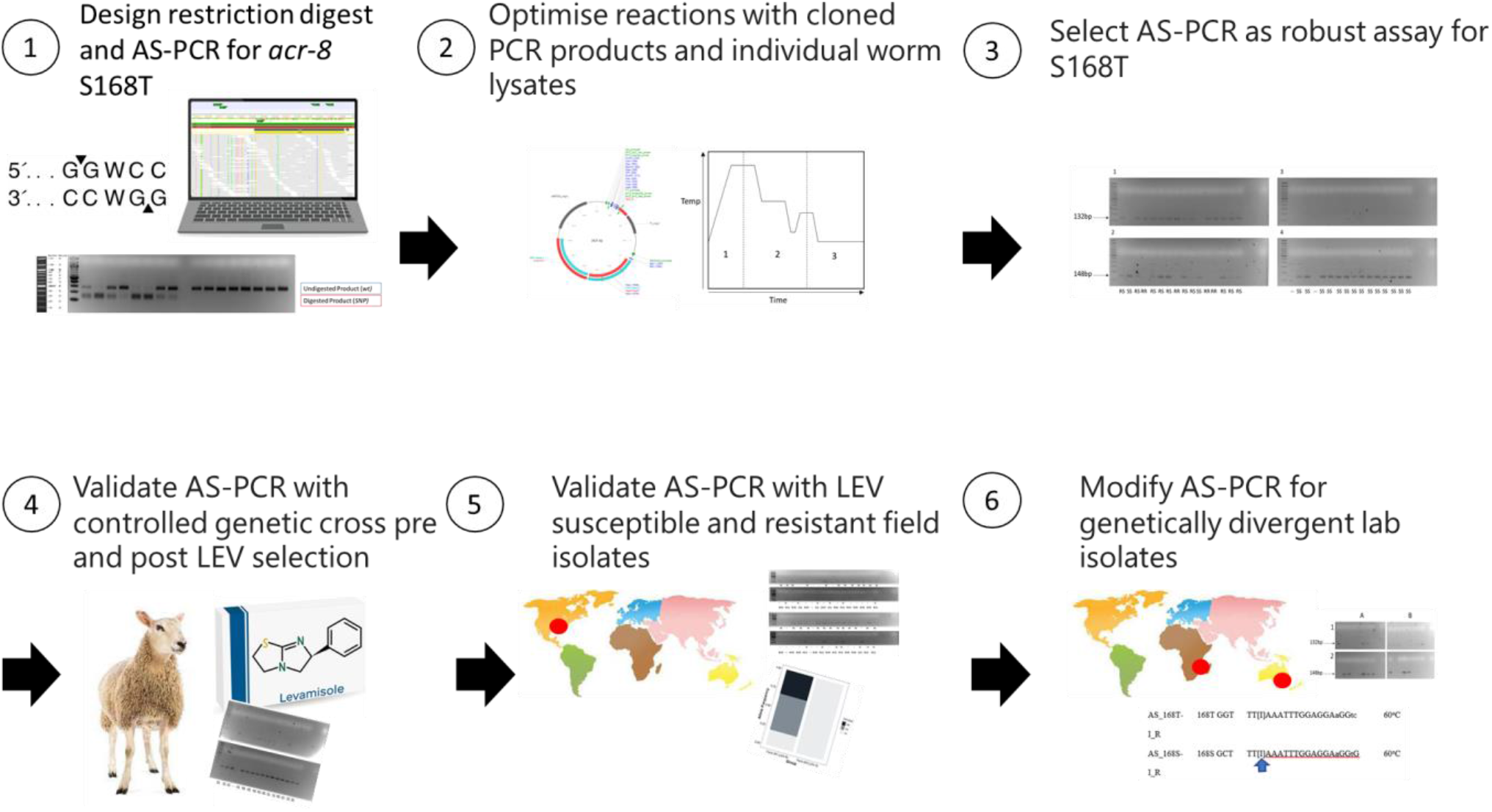
Flow diagram summarising design, establishment, and validation of the AS-PCR for the detection of the S168T variant in multiple *H. contortus* isolates: the LEV susceptible MHco3(ISE), MHco4(WRS), and MHco10(CAVR), and the LEV resistant MHco18(UGA2004) and MHco3/18, and one phenotypically LEV resistant, and one phenotypically LEV susceptible field isolate from the USA.

Allele-specific primers were designed to discriminate between GCT and GGT alleles and PCR conditions were optimised as described in Methods (2.5.4, 2.5.5). These primers were initially used to genotype individual L_3_ from MHco18(UGA2004) and MHco3(ISE) populations. 100% of MHco3(ISE) L_3_ were homozygous for GCT (SS) (n=32) (Figure 5A), while ~96% of MHco3(UGA2004) (n=71) showed at least one GGT (R) allele, of which ~31% were homozygous GGT (RR) and ~65% were heterozygous (RS), with ~4% homozygous GCT (SS) (Figure 5B).

**Figure 5:**
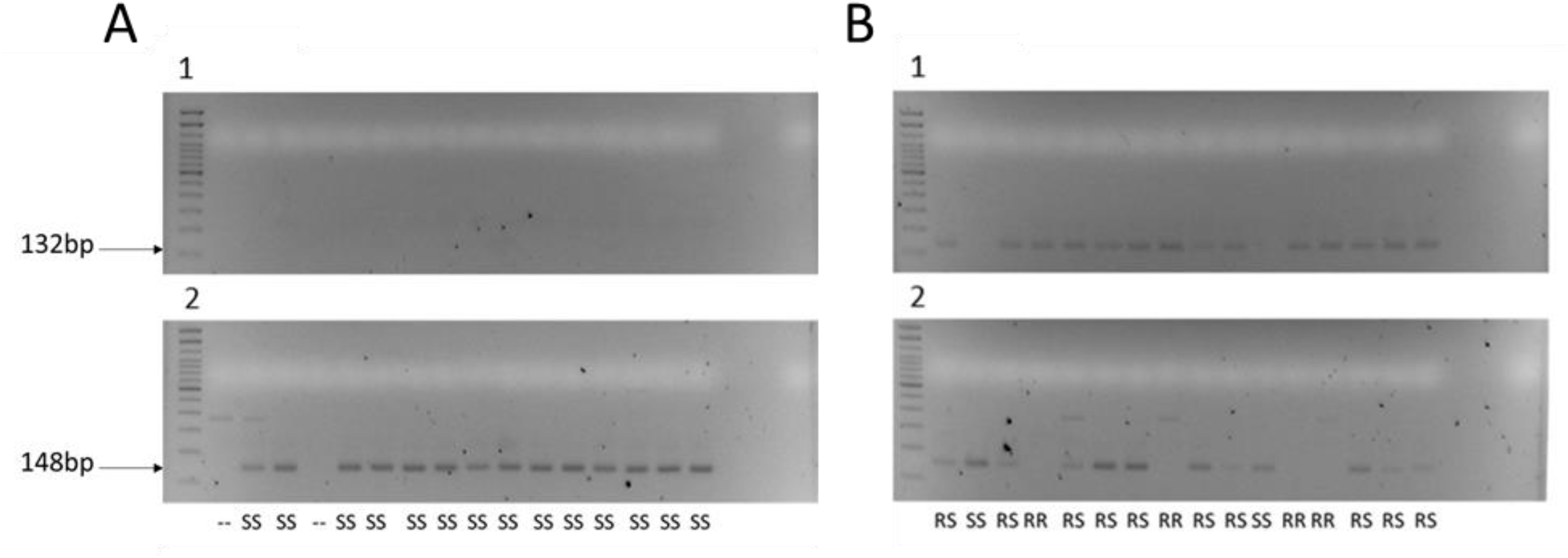
16 LEV resistant MHco18(UGA2004) and 16 LEV susceptible MHco3(ISE) L_3_ were analysed on 2% agarose gel: A: MHco3(ISE): 1: Hco-Exon4-F + Hco-168T-R. 2: Hco-Intron4-F + Hco-168S-R. B: MHco18(UGA2004): 1: Hco-Exon4-F + Hco-168T-R. 2: Hco-Intron4-F + Hco-168S-R. Predicted genotypes RR: homozygous GGT; RS: heterozygous; SS: homozygous GCT. “--” indicates this individual did not amplify by PCR. Only wells that amplified by PCR are counted in the final percentages (Figure 6; Supplementary Table 3).]

We then genotyped F3 larvae of the MHco3/18 cross collected pre- and post-LEV administration. A moderate proportion of GGT alleles were expected in the pre-treatment samples as this population is an admixture of MHco18(UGA2004) and MHco3(ISE) and the RFLP assay previously showed a high proportion of individuals in the untreated population encoded at least one GGT allele (Figure 3; Supplementary Table 2). With AS-PCR, pre-LEV administration, ~58% MHco3/18 showed at least one GGT (R) allele, of which ~9% were RR and 48% were RS, with ~42% SS (n=85). Post-LEV administration, the proportion of individuals with a resistance allele increased: ~75% of MHco3/18 showed at least one GGT (R) allele, of which ~16% were RR and ~58% were RS, with ~25% SS (n=79) (Figure 6; Supplementary Tables 3 and 4). The pre-treatment population was in Hardy-Weinberg equilibrium (HWE) (*χ*^2^ = 0.81, p = 0.29), whereas post treatment the population was not in HWE (*χ*^2^ = 3.26, p = 0.043), with an excess of heterozygotes.

**Figure 6:**
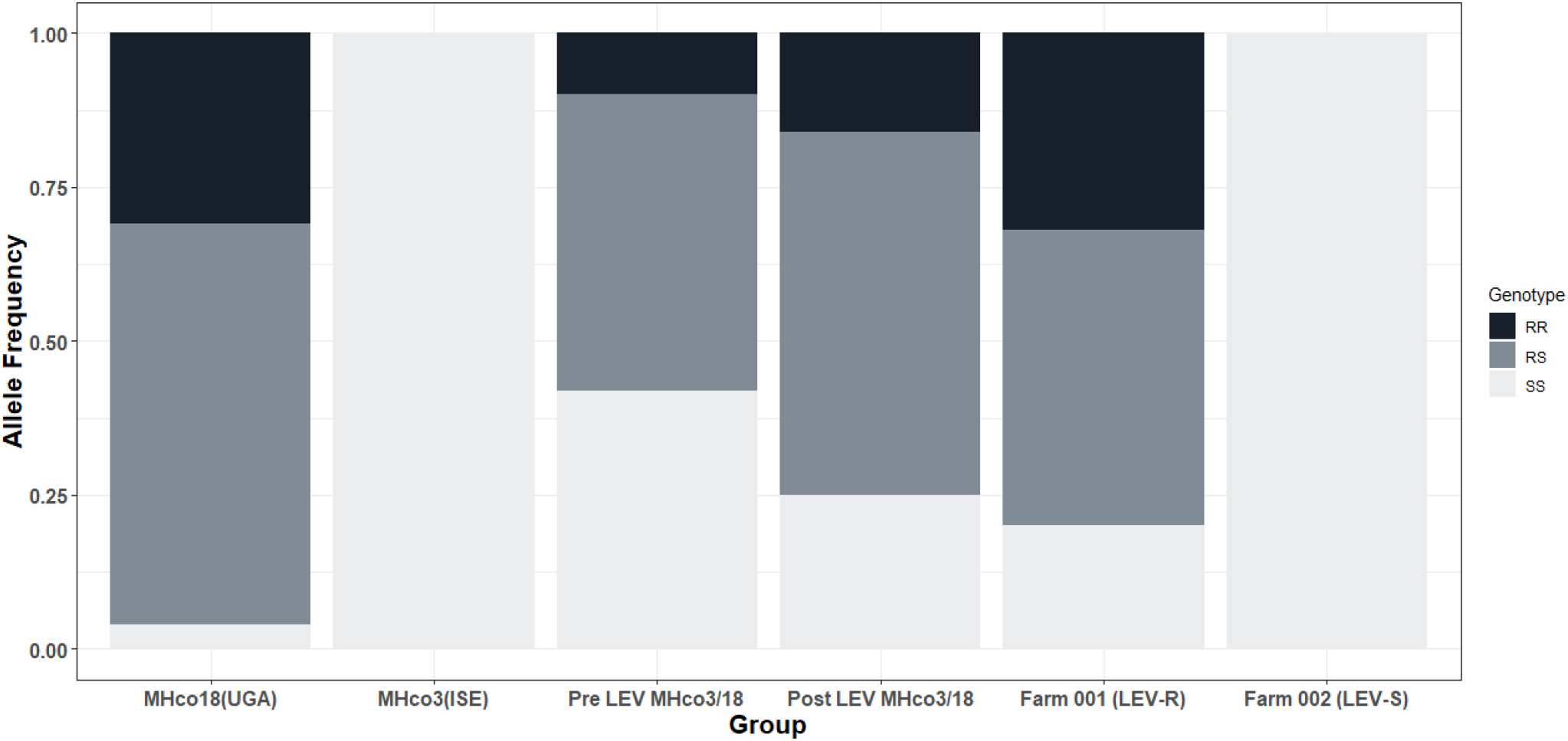
Bar chart showing the proportion of GGT allele detected by AS-PCR in LEV resistant MHco18(UGA2004), LEV susceptible MHco3(ISE), and pre and post single LEV administration MHco3/18, LEV resistant (Farm 001) and LEV susceptible (Farm 002) field populations from the United States of America. Genotypes RR: homozygous GGT; RS: heterozygous; SS: homozygous GCT.

Finally, the AS-PCR was used to genotype *H. contortus* field populations from two farms in the United States of America. DrenchRite® larval development assays predicted Farm 001 to be LEV resistant (EC_50_: 9.36 μM) and Farm 002 to be LEV susceptible (EC_50_: 0.57 μM). 80% of L_3_ from Farm 001 showed at least one GGT (R) allele, of which 32% were RR and 48% were RS, with 20% SS (n=59). In contrast, 100% of individuals from Farm 002 were SS (n=38) (Figure 6; Supplementary Table 3). Farm 002, however, also showed a high percentage of amplification failure (only 59% of 64 wells amplified).

A control PCR targeting a conserved multi-copy locus (ITS2) (Redman et al., 2008) was carried out on the 64 individuals from Farm 002 to determine if the PCR failure in many wells was due to the sample quality of the starting material or polymorphism at the *acr-8* primer binding sites. Following *H. contortus* ITS2 PCR, there was a strong correlation between the band intensity of the two independent PCRs (Supplementary Figure 2). This suggested that the sample quality and/or the concentration of the sample was impacting both PCRs in the same way, indicating degradation of sample material was the most likely explanation, and not genetic variation differentially impacting the two populations.

### 3.4 Deoxyinosine modification of primers optimises S168T GGT allele detection in divergent LEV susceptible isolates

Following optimisation of the AS-PCR in MHco3(ISE) and MHco18(UGA2004), it was tested on two geographically separated LEV susceptible laboratory isolates MHco4(WRS) (Van Wyk et al., 1987) and MHco10(CAVR) (Le Jambre, 1995). These isolates were originally derived from field populations in South Africa and Australia, respectively. The S168T variant (GGT) is not present in pooled WGS sequencing data available for these isolates (Doyle et al., 2019). Initial experiments using the standard AS-PCR primers and conditions (2.5.4) showed no amplification at the previously optimal dilution of 1:20 used for the first-round PCR. The dilution factor was then decreased to 1:8, and although this led to improvements in amplification, it also led to non-specific annealing of the resistant primer (Figure 7A and B). However, using the deoxyinosine substituted primers Hco-168T[I]-R and Hco-168S[I]-R resulted in a marked improvement in both the specificity and sensitivity of the amplification in MHco4(WRS) and MHco10(CAVR) (Figure 7C and D). The deoxyinosine primer showed all individuals assayed to be homozygous for the S168 allele.

**Figure 7:**
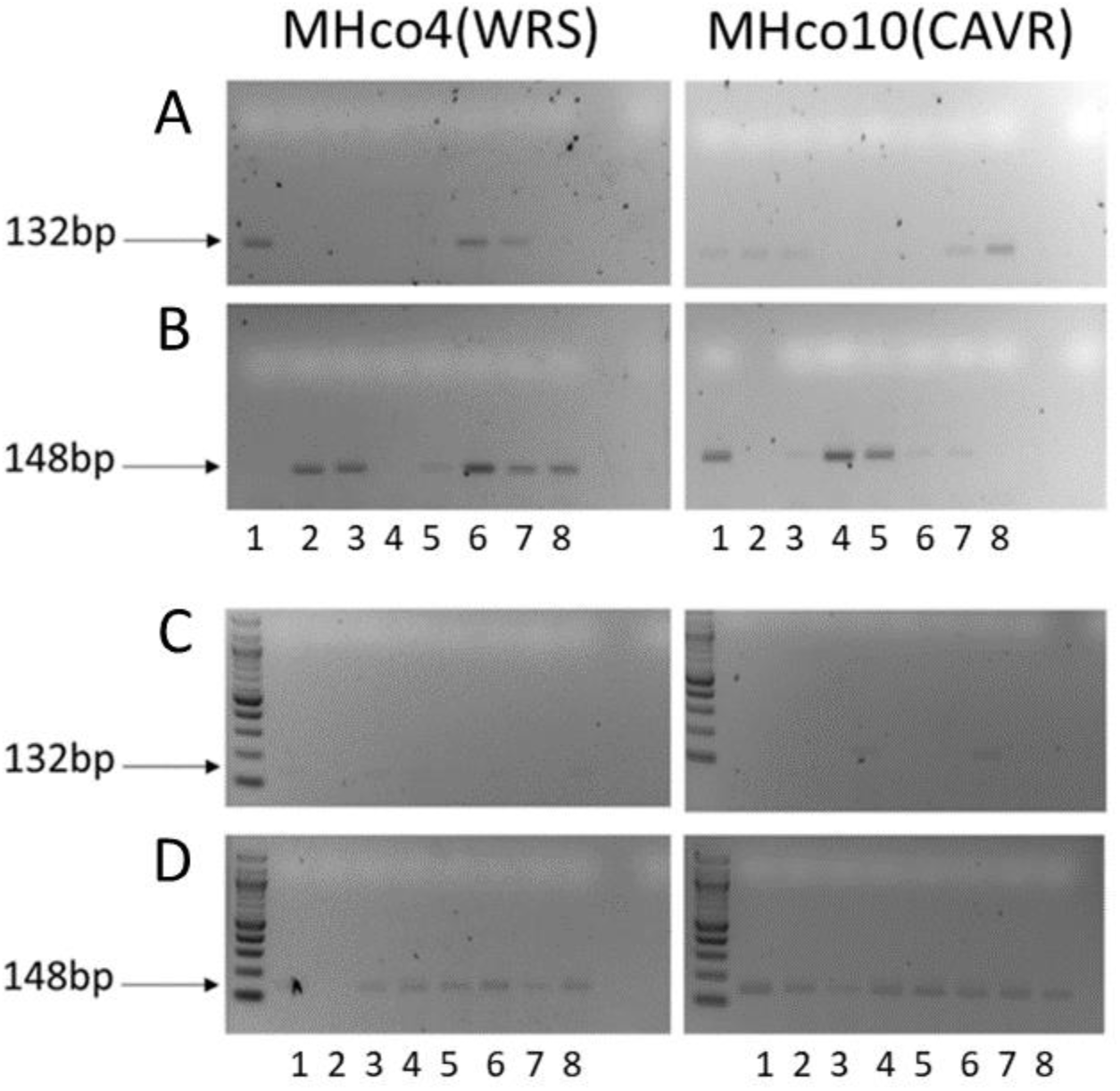
Comparison gel showing difference between unmodified (Hco-168T-R/Hco-168S-R) and deoxyinosine modified (Hco-168T[I]-R/Hco-168S[I]-R) primers on geographically divergent LEV susceptible isolates MHco4(WRS) and MHco10(CAVR): Hco-168T-R/Hco-168S-R primers (panels A and B) 1:8 diluted first round exon 4 PCR products from MHco4(WRS)MHco10(CAVR). Hco-168T[I]-R/Hco-168S[I]-R primers (C and D) 1:20 diluted first-round exon 4 PCR products from MHco4(WRS) and MHco10(CAVR). X axis 1-8: individual L_3_ worms. The same eight individual L_3_ from each isolate were used to compare primer sets.

## 4.1 Discussion

In this study, we sought to characterise putative genetic mechanisms of LEV resistance, with the aim of developing a proof-of-concept diagnostic test that could be used to genotype LEV resistant and susceptible *H. contortus* populations. Proof-of-concept diagnostic assays are considered an essential step towards improving resistance management and surveillance (Kotze et al., 2020), particularly in the face of growing multi-drug resistance (Rose-Vineer et al., 2021). Given the current limitations of the available tests to diagnose LEV resistance in *H. contortus*, we sought to directly address this need by developing an allele-specific PCR.

We undertook extensive single worm genotyping to investigate the presence of the intron 2 deletion. Data from WGS showed that the deletion allele was present within untreated L_3_ of the MHco3/18 genetic cross and increased in frequency following LEV treatment, but this analysis was based on sequencing of pooled L_3_ (Doyle et al, 2022). In the current study, we demonstrated using single worm genotyping that the indel was present and homozygous at a high frequency in both susceptible and resistant isolates. Previous studies have suggested that the high proportion of deletion alleles seen in susceptible populations is due to heterozygous individuals, and thus qPCR could be used as a quantitation-based prediction of resistance in a population (dos Santos et al., 2019). However, this was not supported by our results; given that the majority of individuals in a LEV susceptible population are homozygous for the deletion, it is unlikely to serve as an effective diagnostic marker of LEV resistance. Our results are more consistent with a recent study using droplet digital PCR comparing the presence of the intron 2 deletion in Swedish field-derived and laboratory populations of *H. contortus;* although there was a difference in genotype frequencies between resistant and susceptible isolates, the homozygous deletion was present in a high proportion of susceptible isolates (Baltrušis et al., 2021).

We then sought to characterise the S168T variant, the only non-synonymous SNP in *acr-8* in the MHco18(UGA2004) isolate relative to the fully susceptible MHco3(ISE) isolate (Doyle et al., 2022). Fortuitously, the S168T (GCT/GGT) variant produced as an allele specific restriction site that could be differentiated by an RFLP assay (Figure 3). However, while the RFLP was found to accurately predict the presence of the GGT allele, it was at times difficult to determine if an individual was RR or RS due to the presence and varying intensity of the upper (uncut) fragment. In addition, the requirement for overnight digestion with a restriction enzyme makes the RFLP sub-optimal for a large-scale study or eventual deployment to a diagnostic laboratory.

To overcome these issues, we designed an AS-PCR for quicker and more accurate genotyping of *H. contortus* individuals. The AS-PCR offers marked improvement compared to RFLP, and offers a faster, cheaper, and more accurate assay. AS-PCR eliminated the issue of ambiguous heterozygotes and offers a time to result in under three hours (six hours including crude lysate production). The AS-PCR was sensitive and specific on single L_3_, and we were able to demonstrate a high frequency of the S168T variant in LEV resistant laboratory and field populations, and its absence in LEV susceptible laboratory and field populations. As a proof-of-concept diagnostic assay, the AS-PCR provides the potential for significant improvement on current diagnostic practices for LEV resistance based on the FECRT or other *in vitro* assays (Kotze et al., 2020). In addition, as our assay uses a two-step nested PCR, the *acr-8* exon 4 template generated by the first round PCR could also be integrated into the Nemabiome deep amplicon sequencing system (Avramenko et al., 2019; Melville et al., 2020). This has the added benefit of allowing for increased multifunctionality and improved resistance surveillance for LEV on a larger scale.

Previous work on *H. contortus* suggests that, in some isolates, LEV resistance can be considered a recessive trait (Dobson et al., 1996: Sangster et al., 1998). If LEV resistance is recessive and conferred by the S168T variant alone, we would expect all surviving adults should be RR and all progeny should be RR with no survival of RS or SS genotypes. However, although we observed a significant increase in RR individuals after treatment, we also identified a large proportion of RS individuals in addition to a small proportion of SS larvae. To investigate this observation further we genotyped a small number of F2 adults (n=10) surviving LEV treatment at the S168T locus and found the presence of two SS individuals (data not shown). The presence of fully susceptible adults following LEV treatment suggests either S168T is a linked marker, rather than a mutation conferring resistance, or there are additional resistance mechanisms beyond *acr-8* S168T alone (Neveu et al., 2010; Fauvin et al., 2010; Boulin et al., 2011). Given the conservation of serine at codon 168 across different nematode species and the presence of S168T in LEV resistant *T. circumcincta* (Choi et al., 2017; Doyle et al., 2022), we believe it is likely that S168T is involved in the mechanism of resistance. Furthermore, we have recently identified a second QTL in the MHco3/18 cross following LEV selection containing HCON_00107690 (*lev-1.1*) and HCON_00107700 (*lev-1.2*) on Chromosome IV (Doyle et al., 2022), which is suggestive of additional mechanisms of resistance. *lev-1* has previously been implicated in LEV resistance in *C. elegans* (Fleming et al., 1997; Qian et al., 2008). However, its role in LEV resistance in *H. contortus* remains unclear, as it is thought to lack a signal peptide for membrane insertion, and experiments in *Xenopus* oocytes showed that *H. contortus* LEV-1 was not required for a functional reconstituted receptor (Boulin et al., 2011). Further work is required to understand the relative contribution of mutations in *acr-8* and at the *lev-1* locus, as well as in other commonly implicated loci, in the LEV resistance phenotype.

A limitation of a PCR-based diagnostic is the potential for genetic variation in the primer binding sites to interfere with the primer binding and therefore assay efficiency. This is a legitimate concern, given that populations of *H. contortus* are known to be highly genetically diverse throughout the world (Yin et al., 2013; Sallé et al., 2019). To investigate the possibility of developing an assay for S168T in divergent populations, the AS-PCR was tested on two LEV susceptible isolates, MHco4(WRS) and MHco10(CAVR), from South Africa and Australia respectively. We initially found that the AS-PCR performed poorly on these isolates, and confirmed via ITS2 species identification PCR, that this was likely due to sequence differences rather than availability of starting template. As simply decreasing the dilution factor in these isolates led to non-specific amplification, deoxyinosine [I] bases were introduced in the primers to accommodate two synonymous SNPs that were identified at the primer bind site. [I] base pairing allows for “universal” pairing at base positions likely to exhibit polymorphism, albeit with a preferential order for pairing (I-C>I-A>I-G/I-T) (Case-Green and Southern, 1994). This follows an approach used to develop a universal (i.e. detecting all four serotypes) Dengue Virus PCR primer set (Wang et al., 2000), a pathogen that also displays very high levels of sequence polymorphism. The introduction of a single [I] base (Hco-168T[I]-R/Hco-168S[I]-R) at the site of the most frequent SNP led to a marked increase in both specificity and sensitivity of the assay, restoring amplification at 1:20 dilution. The consistency of the amplification was also markedly improved. The introduction of a second [I] base (Hco-168T[II]-R/Hco-168S[II]-R) however, compromised the primer binding under the current reaction conditions at all dilutions tested (data not shown). Thus, further optimisation will be necessary to overcome the issues of non-specific amplification and produce a truly global *H. contortus* LEV resistance assay primer set. Alternatively, it may be necessary to develop region specific primer sets for optimal diagnostic performance, an undertaking that would be facilitated by the global diversity database of *H. contortus* (Sallé et al., 2019).

## 4.2 Conclusion

In conclusion, we investigated two mutations at the *acr-8* locus proposed to be associated with LEV-resistance in *H. contortus*. Of these, only the SNP marker S168T consistently discriminated between susceptible and resistant L_3_ of all populations tested by single worm genotyping. While it is likely that LEV resistance is multigenic, our data further implicates S168T as a major diagnostic marker. We developed an optimised AS-PCR for the detection of this SNP in *H. contortus* and show that the modification of a primer set with an [I] base presents a potential solution to dealing with genetically divergent populations. This proof-of-concept molecular assay provides the starting point for a sensitive diagnostic test for LEV resistance in global populations of *H. contortus*.

## Acknowledgements

This work was funded by a James Herriot Scholarship (Glasgow University Vet Fund) and a Biotechnology and Biological Sciences Research Council (BBSRC) strategic Lola [BB/M003949]. SRD is supported by a UKRI Future Leaders Fellowship [MR/T020733/1] and RL is supported by a Wellcome Clinical Research Career Development Fellowship [216614/Z/19/Z]. DB and AM are supported by the Scottish Government’s Rural and Environment Science and Analytical Services (RESAS) division. This research was funded in whole, or in part, by the Wellcome Trust. For the purpose of open access, the author has applied a CC BY public copyright licence to any Author Accepted Manuscript version arising from this submission.

