## Supplementary Figures for "Allele specific PCR for a major marker of levamisole resistance in *Haemonchus contortus*"

c: Moredun Research Insititue, Penicuik, Scotland

d: St. George's University, Grenada

e: Department of Infectious Diseases, College of Veterinary Medicine, University of Georgia, USA

f: Institut National de la Recherche Agronomique, Nouzilly, France

g: Institute of Biodiversity, Animal Health, & Comparative Medicine, University of Glasgow, Glasgow, Scotland

This PDF file includes:

Supplementary Figure 1

Supplementary Figure 2

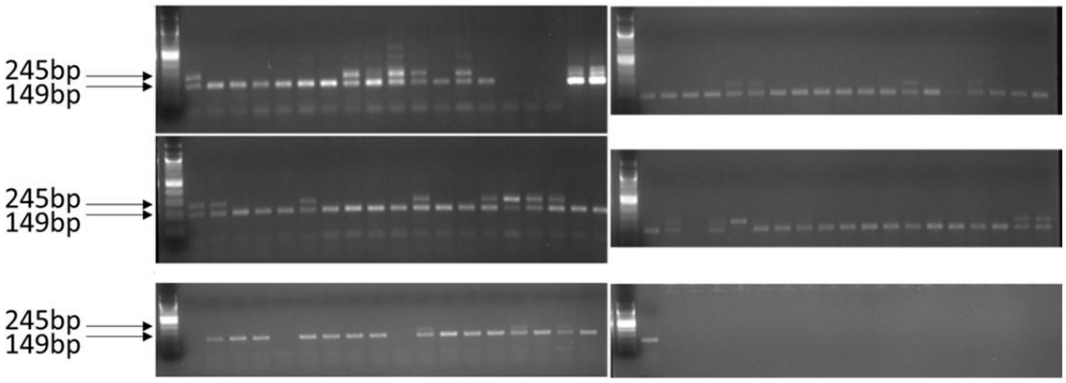


[Supplementary Figure 1: Example gel showing large sample (n=94) of MHco3(ISE) using size discrimination Indel PCR using primers Hco-Indel-F and Hco-Indel-R.]

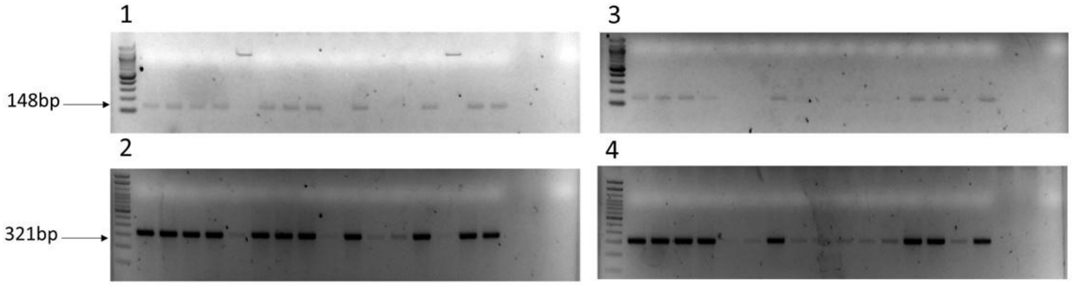


[Supplementary Figure 2: Example gel showing amplification of Farm 002 by AS-PCR (1 and 3; Hco-168S-R (sensitive) primer; 148 bp band) and ITS2 speciation PCR (2 and 4; 321 bp band) and 100% concordance between poor amplification by AS-PCR and ITS2 PCR indicating a lack of genetic material is responsible for failure of those wells to amplify.]
